# Single-cell multiomic profiling reveals lineage plasticity in pediatric B-lineage Acute Lymphoblastic Leukemia during the early phase of treatment

**DOI:** 10.64898/2026.06.15.730052

**Authors:** Giulia Gomiero, Alberto Peloso, Elena Varotto, Pamela Scarparo, Martina Volgger, Chiara Frasson, Alice Cani, Grazia Fazio, Giovanni Cazzaniga, Fraia Melchionda, Anna Maria Testi, Franco Locatelli, Carmelo Rizzari, Alessandra Biffi, Michael N. Dworzak, Silvia Bresolin, Barbara Buldini

## Abstract

The biological bases of transient myelomonocytic switch (mmSW) during induction therapy in B-cell precursor acute lymphoblastic leukemia (BCP-ALL) remains largely undefined. Here we integrated single-cell transcriptomic and surface marker profiling with genomic and DNA methylation analyses of pediatric BCP-ALLs, including matched diagnosis (Dx) and Day+15 samples from mmSW-positive (mmSWpos) and mmSW-negative cases. At Dx, mmSWpos leukemia samples were enriched for hematopoietic stem/progenitor-like cell subpopulations. Trajectory and entropy analyses identified a pre-existing “fate-uncertain” cell compartment co-expressing both lymphoid and myeloid programs. Longitudinal single-cell data showed that, in mmSWpos samples, this population undergoes complete transdifferentiation. mmSWpos are enriched for Ras pathway and chromatin regulation mutations and display a specific DNA hypermethylation pattern at Dx. These findings indicate that transient mmSW arises from intrinsic leukemic plasticity, in which immature transcriptomic and distinct epigenetic states at diagnosis enable lineage switching under the pressure of ALL treatment.

## MAIN

Advances in omics technologies allowed the definition of distinct B-cell precursor acute lymphoblastic leukemia (BCP-ALL) subtypes characterized by specific genetic, transcriptomic, and immunophenotypic features^1^. Genetic abnormalities associated with lineage infidelity in acute leukemia have progressively expanded and have been incorporated in the latest version of the WHO Classification^2^. Immunophenotypic features of acute leukemia may evolve during treatment or at relapse, a phenomenon well known in *BCR::ABL1* fusion or *KMT2A* rearrangements bearing cases^3–6^. Over the last decade, few BCP-ALL subgroups without *KMT2A* rearrangements were shown to exhibit a change of blasts immunophenotype, described as myelomonocytic switch (mmSW), in the first phase of induction therapy^7^. Different mmSW patterns in induction therapy have been recognized in three distinct genetic subtypes, namely *DUX4*-rearranged (*DUX4*-r), *ZNF384*-rearranged (*ZNF384*-r), and PAX5-P80R-mutated^8^. In 2024, the AIEOP BFM Flow Network fully dissected the immunophenotypic behavior of a peculiar subtype of BCP-ALL characterized by the aberrant expression of CD371 marker at diagnosis and a transient mmSW during the early induction phase in up to 60% of cases, associated with *DUX4* rearrangement^9,10^. This phenomenon is defined by the appearance of a population of blasts showing a down-regulation of CD19 and up-regulation of myeloid markers, in addition to blasts with the same immunophenotype of diagnosis^8,10^. mmSW is efficaciously controlled by ALL-oriented therapy without requiring any shift to acute myeloid leukemia-oriented treatment and the prognosis is favorable in most patients.

However, the mechanisms underlying this transient mmSW remain to be elucidated. Different hypotheses have been proposed to explain the phenomenon: (i) occurrence of a direct trans-differentiation or indirect redifferentiation of an already committed B-cell progenitor; (ii) presence of a pluripotent stem cell or a bipotent B-myeloid precursor already present at diagnosis that could be reprogrammed; (iii) selection of a subclone already present at diagnosis^3^.

In this study, we investigated by a multiomic approach the immunophenotypic plasticity of pediatric BCP-ALL cases undergoing mmSW in the first phase of induction therapy, revealing the presence of a highly dynamic leukemia stem cell population prone to transdifferentiate toward myeloid lineage.

## RESULTS

### mmSWpos BCP-ALL blast population is transcriptionally distinct at diagnosis

We applied a multiomic approach to primary bone marrow (BM) samples from pediatric BCP-ALL patients who underwent (or not) mmSW to systematically characterize this phenomenon (Fig. 1a). We generated scRNA-seq and surface markers (AbSeq) data from samples collected at diagnosis from 11 pediatric BCP-ALL with distinct genetic backgrounds (9 *DUX4*-rearranged, one *ZNF384*-rearranged, and one with *ETV6::RUNX1* fusion, Supplementary Table 1) and different propensity to develop transient mmSW during induction therapy. Genetic subtypes were initially assigned by fusion detection and gene expression-based classifiers on bulk RNA-seq and subsequently confirmed in pseudobulk scRNA-seq profiles (Supplementary Table 1). MmSW was detected by Multiparametric Flow Cytometry (MFC) at Day+15 of induction therapy (D15), as previously described^10^. Patients with evidence of mmSW at D15 were defined as mmSWpos (n=8), whereas those with B-lineage leukemia blasts with no immunophenotypic change at the same time point were defined as mmSWneg (n=3) (Supplementary Table 1).

**Fig. 1:**
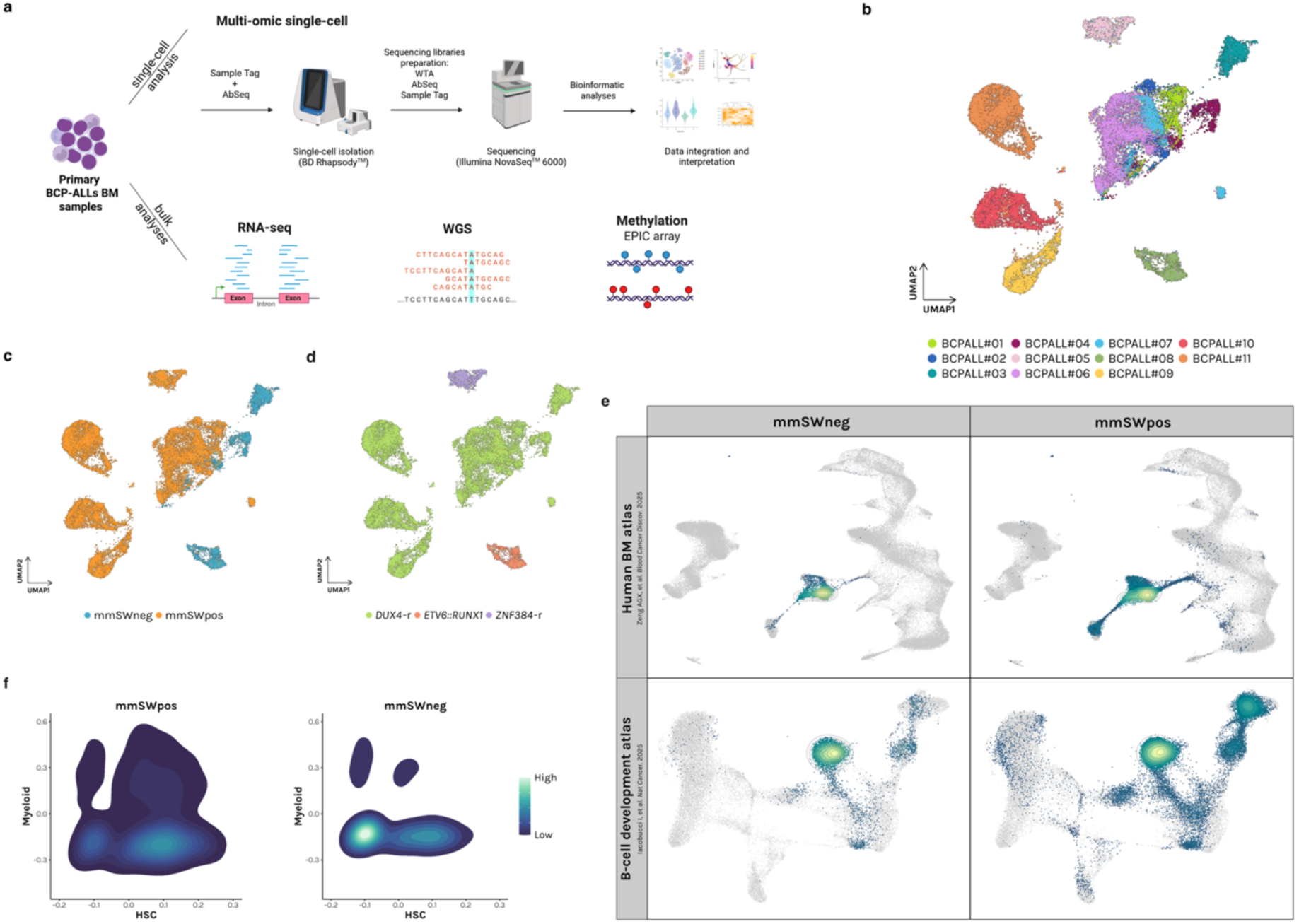
A distinct transcriptional state defines mmSWpos BCP-ALLs blasts at diagnosis. **a**, Schematic representation of the experimental workflow used for the characterization of BCP-ALLs BM samples by single-cell and bulk multi-omics. The figure was created with biorender.com. **b-d,** UMAP embeddings of 31,597 leukemic cells from 11 BCP-ALL BM samples (n = 3 mmSWneg, n = 8 mmSWpos) at diagnosis. Colors represent different patients in (**b**), while in (**c**) mmSWneg and mmSWpos are colored in blue and orange, respectively. In (**d**) colors represent genetic subtypes. **e,** Projection of blasts at diagnosis (n = 11 BCP-ALLs) onto reference atlases of human BM (Zeng AGX et al. *Blood Cancer Discov*. 2025) and B-cell development (Iacobucci I et al. *Nat Cancer*. 2025). Density plots indicate preferential localization of projected cells along the differentiative trajectories and suggest their maturative stage. **f,** Density distribution of mmSWpos (n = 8) and mmSWneg (n = 3) blasts along the hematopoietic stem cell (HSC) and myeloid axes, showing subgroup-specific transcriptional priming within the developmental landscape. Priming scores were computed using myeloid signature from Pellin D et al. *Nat Commun.* 2019 and HSC signature from Hao et al. *Cell.* 2021.

We acquired a total of 35,127 high-quality cells with joint whole transcriptome and 14-plex AbSeq profiles, obtaining a high-resolution BM picture of BCP-ALL (Extended Data Fig.1a). AbSeq were selected based on the antibody panels routinely used for immunophenotyping at diagnosis and minimal/measurable residual disease detection^11,12^. To discriminate leukemic blasts from non-malignant BM cells, we annotated cell populations by integrating transcriptomic and AbSeq data. Cells from different patients that clustered together, annotated as mature B-, myelo-erythroid and T-cells and being CD20^+^/CD45^+^, CD33^+/-^/CD45^+/-^and CD2^+^/CD45^+^ respectively, were excluded from downstream analyses as components of the non-malignant BM compartment (Extended Data Fig.1b-c). The expression of canonical lineage-defining genes confirmed the non-malignant identity of these populations (Extended Data Fig.1d-e). In addition, inferCNV analysis allowed us to further distinguish blasts from the non-malignant clusters at a single-cell level (Extended Data Fig.1f).

Blast cells exhibited variable expression of CD19, CD34, CD38, CD10, and CD371 (in CD371^+^ BCP-ALLs) at the protein level with AbSeq, consistent with MFC at diagnosis (Extended Data Fig.2 and Supplementary Table 2).

Single-cell transcriptomic clustering revealed that blasts from different patients, including those within the same genetic subtype, segregated into distinct clusters, highlighting a high heterogeneity in BCP-ALL, in line with recent reports.^13,14^ (Fig. 1b-d). Notably, blasts from mmSWpos patients grouped separately from mmSWneg ones, underlining that these two subgroups are already transcriptionally distinct at diagnosis (Fig. 1c).

To delineate the ontogeny of blast cells in mmSWpos and mmSWneg BCP-ALLs at diagnosis, we projected single-cell RNA-seq data onto a healthy reference^15^ that encompasses all BM developmental stages (Extended Data Fig.3a). Blasts predominantly mapped along the B-cell development trajectory (Fig. 1e, upper panel). Therefore, to provide a fine-mapping projection of the differentiative stages along B-cell lineage, we leveraged a second, higher-resolution B-cell ontogeny reference^13^ (Extended Data Fig.3b). mmSWneg samples showed an exclusive abundance of cells inside the “B-cell progenitor” stage (i.e., Pro-B VDJ). Conversely, mmSWpos blasts displayed a more heterogeneous distribution across the entire B-cell differentiation, with representation at nearly all maturation stages and a notable enrichment in the hematopoietic stem cells (HSC)/ multipotent progenitors (MPP) compartments (Fig. 1e lower panel and Extended Data Fig.3c-d). Consistently, mmSWpos blasts at diagnosis exhibited an HSCs-like^16^ transcriptional priming and a tendency to myeloid commitment^17^ (Fig. 1f).

Our findings highlight that blast cells of patients undergoing a mmSW during the early stage of induction therapy are transcriptionally distinct at diagnosis, with an enhanced stem– and myeloid-priming, suggestive of increased immunophenotypic plasticity.

### mmSWpos BCP-ALLs are characterized at diagnosis by a subpopulation prone to transdifferentiation towards myelomonocytic lineage

Next, we aimed at depicting the molecular mechanisms of immunophenotypic plasticity during the early phase of induction therapy. Thus, using the same multiomic single-cell approach, we were able to profile 6 BM sample pairs collected at diagnosis (Dx) and at D15: 5 mmSWpos (BCPALL#06, BCPALL#07, BCPALL#09, BCPALL#10, BCPALL#11) and one mmSWneg (BCPALL#08) (Extended Data Fig.4a-b and Supplementary Table 1, 3). We leveraged AbSeq and lineage-specific marker genes as defined at Dx to annotate blasts and exclude non-malignant B, myelo-erythroid, and T cells from downstream analyses (Extended Data Fig.4c-d). Patient-specific distribution of blasts confirmed the intrinsic heterogeneity of the disease both at Dx and D15 (Fig. 2a). Notably, while D15 blasts in the mmSWneg case largely overlapped with Dx, D15 blasts in mmSWpos patients clustered distinctly from those characterized at diagnosis (Fig. 2a-b), indicating the occurrence of a clear transcriptomic shift. Up-regulation of myeloid markers (i.e. CD11b, CD14, CD33) and simultaneous down-regulation of lymphoid markers confirmed the immunophenotypic data previously described by MFC at D15.^10^ (Extended Data Fig.4e).

**Fig. 2:**
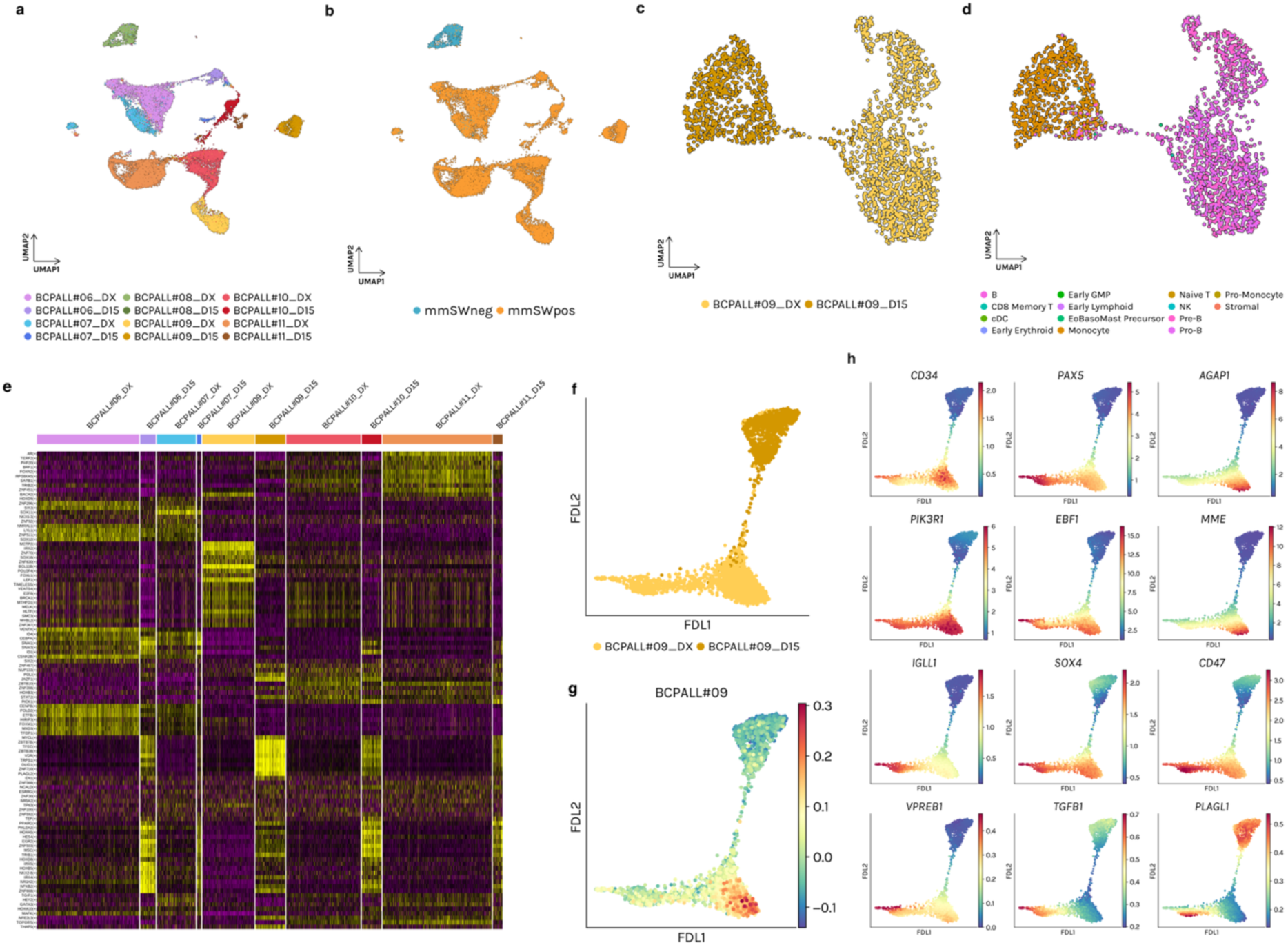
Single-cell transcriptomic landscape and differentiation trajectories of BCP-ALLs at diagnosis and D15. **a-b**, UMAP plot of 28,105 blast cells from 12 samples collected at Dx and D15. Cells are colored by sample identity in (**a**) and for mmSW status in (**b**). **c,** UMAP of coupled Dx-D15 samples from a representative patient (BCPALL#09), highlighting the distribution of cells between the two timepoints. Different shades of yellow indicate timepoints. **d,** Cell-type labelling based on projection onto healthy BM reference in a representative patient (BCPALL#09). Cells are colored by BM subpopulation. **e,** Heatmap of regulon activity inferred by pySCENIC, showing activity scores across single cells from couples Dx and D15 samples. Rows represent regulons, while columns represent individual cells, grouped by sample. **f,** Force-directed layout (FDL) embeddings of BCPALL#09 revealing a continuous transcriptional trajectory linking Dx and D15 populations. Colors show different timepoints. **g,** *DUX4*-r signature score projected onto FDL, showing a gradient decrease towards D15. Color gradient represent signature score. **h,** MAGIC-denoised gene expression values of markers from B-lineage, immature cells, and genes previously shown to be upregulated in patients undergoing mmSW. MAGIC-imputed gene expression scores are represented as color gradient.

To account for the inter-patient’s high heterogeneity, we performed clustering for each Dx-D15 pair separately (Fig. 2c and Extended Data Fig.5a-b). Single cells from each pair were then projected onto the human whole BM atlas^15^, as done for Dx samples. In mmSWpos samples at D15 we observed a marked depletion of B-cell populations in favor of myelomonocytic cells, demonstrating that the mmSW observed by MFC is also mirrored at the transcriptomic level (Fig. 2d and Extended Data Fig.5a-b). In contrast, D15 cells from the mmSWneg patient (BCPALL#08) were annotated as B-cell progenitors (pro– and pre-B), indicating the absence of a transcriptional lineage reprogramming (Extended Data Fig.5b). To dissect the regulatory programs driving the Dx–D15 transition, we applied pySCENIC to compare regulon activity between the two time points (Fig. 2e). mmSWpos samples exhibited D15-specific activation of shared gene regulatory networks (GRNs). Interestingly, key regulators of HSC– and myeloid-associated programs showed different activities along the Dx-D15 axis (Extended Data Fig.5c-d).

We reconstructed cell-state transitions along the Dx-D15 axis and inferred the cellular origin underlying the transient mmSW by applying force-directed layout (FDL) to single-cell data (Fig. 2f and Extended Data Fig.6a). Thus, evolutionary trajectory between the two timepoints is emphasized, allowing to better map the blast plasticity during mmSW. To further characterize the transcriptional reprogramming, we computed single-cell signature scores reflecting genomic subtype-specific programs. The *DUX4*-r signature^18^ showed a bimodal distribution at Dx and a progressive decline by D15 in mmSWpos, supporting a complete transdifferentiation process of blasts during induction therapy (Fig. 2g and Extended Data Fig.6b). Moreover, diffusion map modeling on FDL-embedded cells confirmed the bimodal distribution of gene expression in mmSWpos blasts at Dx. In detail, one blast subpopulation expressed B-cell commitment genes such as *EBF1, PAX5,* and *MME*, whereas the second subpopulation showed higher expression of myeloid and stemness regulators, including *TGFB1*, *IGLL1*, *SOX4,* and *VPREB1*^14,19^. We also identified a subset of cells co-expressing both lymphoid and myeloid programs at Dx, consistent with a “fate-uncertain” state. Interestingly, *PLAGL1* was upregulated in this latter subpopulation as in part of D15 blasts (Fig. 2h and Extended Data Fig.6c-f). These observations highlight intrinsic heterogeneity and transcriptional instability of mmSWpos blasts at diagnosis.

### Pseudotime ordering and cell entropy analysis revealed a high dynamicity and different timing of mmSW progression

Given our identification of transcriptional heterogeneity in mmSWpos samples, we next leveraged pseudotime inference to trace the differentiation trajectories underlying blast evolution between Dx and D15 (Fig. 3a and Extended Data Fig.7a-d). Pseudotime ordering revealed a highly entropic cell population spanning Dx and D15 in all mmSWpos pairs (Fig. 3b and Extended Data Fig.7a-d), reflecting rapid transcriptional remodeling along this axis. This feature was also confirmed by the application of Mellon^20^, showing that the bridging population is characterized by both low density and elevated entropy (Fig. 3c and Extended Data Fig.7e). Interestingly, the “fate-uncertain” blast subpopulation exhibited moderately high entropy, supporting the hypothesis that these cells are transcriptionally unstable and prone to dynamic state transitions. Cell entropy analysis combined with gene expression modeling delineated two differentiation branches: a ‘B cell’ branch and a ‘Mono’ (monocytic) branch (Fig. 3d and Extended Data Fig.7f-i). Cells following the ‘B cell’ branch mapped predominantly within Dx samples, retained a precursor B-cell phenotype, maintaining high expression of markers such as *MME* and *RAG1* across pseudotime. In contrast, cells progressing along the ‘Mono’ branch showed a gradual acquisition of monocytic features, marked by increasing expression of *CEBPD* and *CD33* and the progressive loss of B-lineage markers (Fig. 3e and Extended Data Fig.7j-m).

**Fig. 3:**
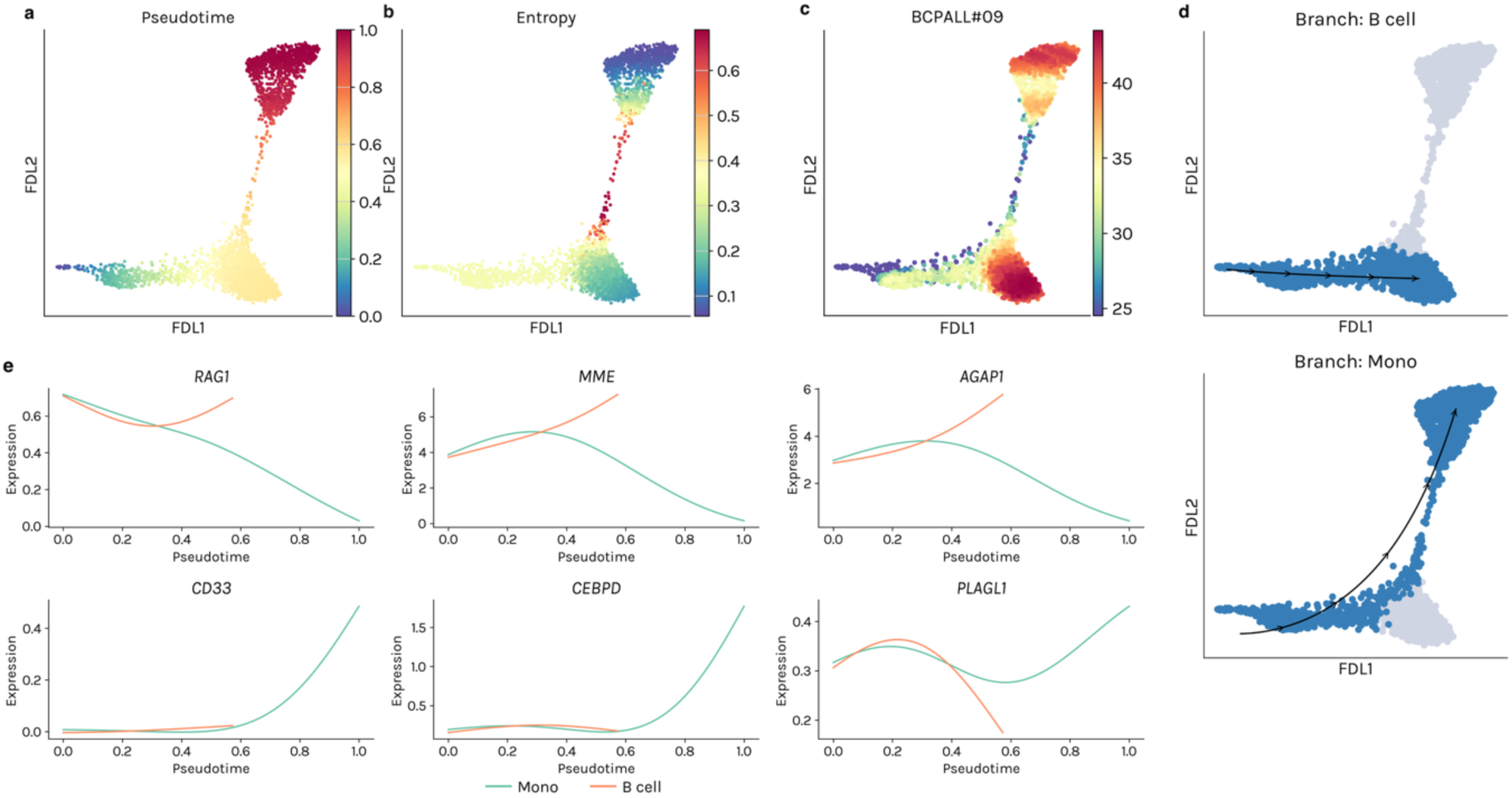
Single-cell pseudotime and trajectory inference identify two differentiation programs in the Dx-D15 axis. **a**, FDL embedding of single cells colored by inferred pseudotime of BCPALL#09, showing a continuous developmental trajectory originating from a common progenitor state and diverging toward two terminal cell fates. Color gradient indicated pseudotime, ranging from 0 to 1 **b,** Palantir cellular entropy mapped onto FDL of BCPALL#09, highlighting regions of higher transcriptional plasticity at early states and reduced entropy along more differentiated trajectories. Color indicates entropy values. **c,** Cell-state densities (clipped) estimated by Mellon across the embedding. Color gradient is indicative of density scores. **d,** Branch assignment and inferred lineage dynamics for the two trajectories defined in BCPALL#09. Arrows indicate progression toward terminal ‘B cell’ (top) and ‘Mono’ (bottom) states. **e,** Gene expression trends along pseudotime in BCPALL#09. Green and orange lines indicate progression toward ‘Mono’ and ‘B cell’ trajectories, respectively.

We observed two distinct patterns of cell state transition from Dx to D15. A progressive increase in entropy, peaking in a transitory intermediate cell state, was detected in a group of patients (BCPALL#07, BCPALL#09, BCPALL#10, BCPALL#11), consistent with an ongoing reprogramming process (Fig. 3d and Extended Data Fig.7g-i). By contrast, BCPALL#06 lacked a gradual entropy increase; instead, entropy rose sharply at D15, indicating a “jump” transition. (Extended Data Fig.7f). Notably, BCPALL#11 displayed a distinctive profile, with most blasts already exhibiting high entropy at Dx, consistent with an expanded “fate-uncertain” compartment and an early propensity towards transcriptional reprogramming (Extended Data Fig.7d-i).

Collectively, these data confirm the vulnerability of mmSWpos patients to a transdifferentiation towards the myelomonocytic lineage during therapy. Although Dx-D15 pairs reveal patient-specific dynamics and different timing of cell-state transitions, they converge on a shared mechanistic framework in which mmSW is driven by the lineage switch of a “fate-uncertain”, transcriptionally unstable blast population.

### mmSWpos BCP-ALLs show a distinct mutational and epigenetic landscape at diagnosis

We next asked whether genomic and epigenomic features of BCP-ALL patients at diagnosis could account for the mmSW observed during induction therapy. We first integrated whole-genome sequencing (WGS) and bulk RNA-seq data to depict the mutational landscape of 21 BCP-ALL patients, 6 mmSWneg and 15 mmSWpos (Supplementary Table 1). Single nucleotide variants in key genes involved in Ras signaling pathway were detected in 38% of patients. Of note, 88% of them are mmSWpos, supporting their myeloid transcriptional priming. In addition, 53% of mmSWpos patients carried variants in genes associated with epigenetic regulation and chromatin remodeling (Fig. 4a, Extended Data Fig.8 and Supplementary Table 5-6).

**Fig. 4:**
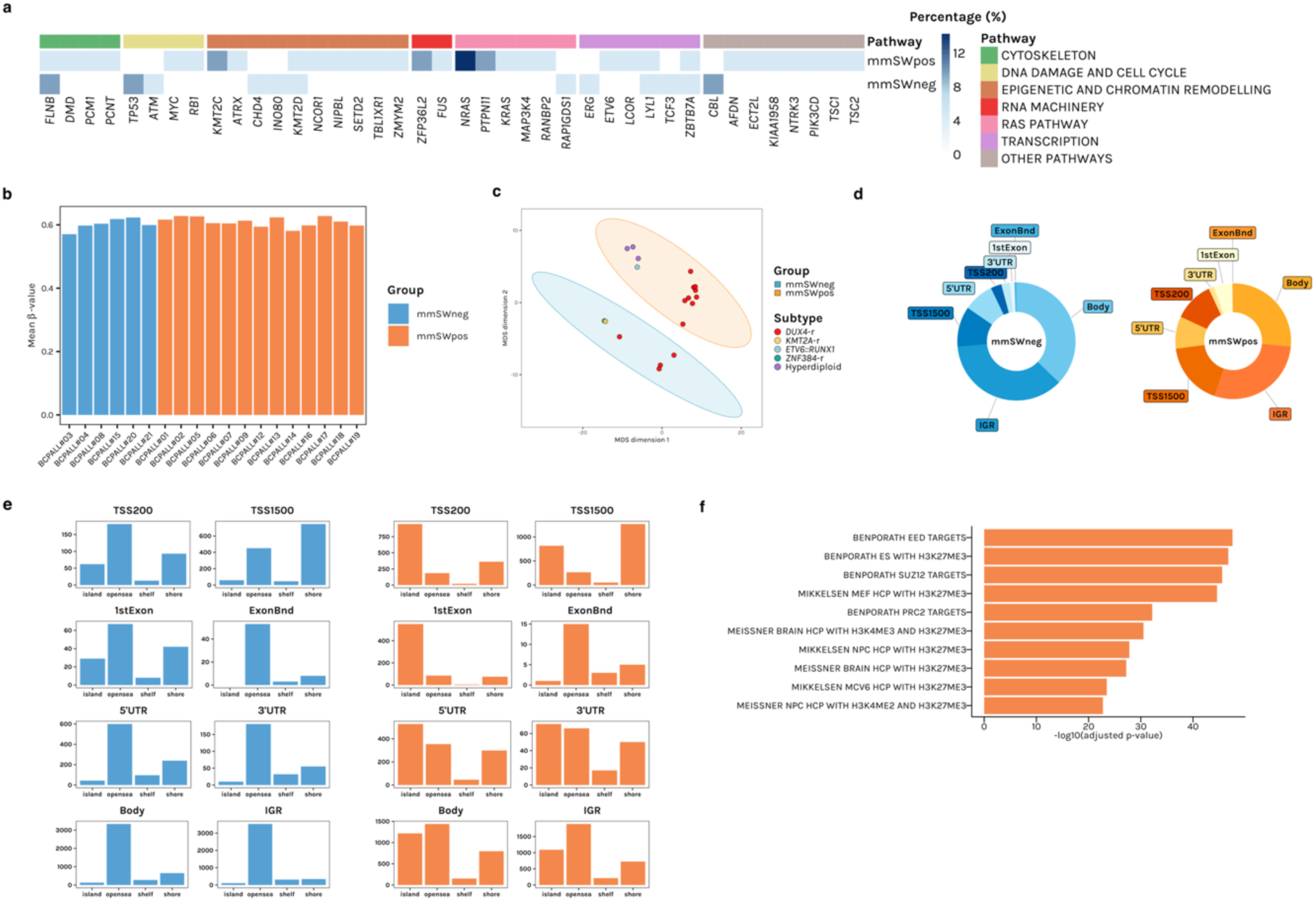
Mutational and epigenetic landscapes in mmSWpos BCP-ALLs at diagnosis. **a**, Heatmap showing recurrently mutated genes across mmSWneg (n = 6) and mmSWpos (n = 15) patients, grouped by functional pathways. Color gradient represents the percentage of patients with a mutation in each gene, calculated across all patients. **b,** Mean β-values were compared between mmSWneg (n = 6) and mmSWpos (n = 13) samples using a two-sided Welch’s t-test (P value = 0.4023). mmSWneg and mmSWpos are shown in blue and orange, respectively. **c,** Multidimensional scaling (MDS) plot based on the 1,000 most variable probes shows a separation between mmSWneg and mmSWpos samples, with a high contribution given also by the genetic subtype. Ellipses define mmSW subgroups, with mmSWneg depicted in blue (n = 6) and mmSWpos in orange (n = 13), indicating 95% of confidence interval. Colors indicate the genetic subtype. **d,** Genomic distribution of differentially methylated probes (DMPs) in mmSWneg (n = 6) and mmSWpos (n = 13) patients. TSS200 and TSS1500 denote regions within 200 bp and 1,500 bp upstream of the transcription starting site, while IGR indicates intergenic regions. **e,** Region context of subgroup-specific DMPs stratified by genomic annotation and CpG context (island, shore, shelf and open sea), depicting the genetic landscapes of methylation in mmSWneg (blue) and mmSWpos (orange) BCP-ALLs. **f,** Gene set enrichment analysis (GSEA) of mmSWpos-associated DMPs. Top 10 enriched gene sets are shown, ranked by –log10 adjusted P value.

To investigate the role of epigenetics in mmSW, we profiled DNA methylation of 19 BM samples collected from patients at diagnosis (6 mmSWneg and 13 mmSWpos). No significant differences (P value = 0.4023) in global mean methylation levels were observed between the mmSWneg and mmSWpos subgroups (Fig. 4b). However, unsupervised analysis of the 1,000 most variable probes revealed that BCP-ALLs clustered differently according to genetic subtypes, with *DUX4*-r patients clearly separated from other subtypes. Despite the contribution of genetics, a clear separation between mmSWpos and mmSWneg was observed, highlighting distinct methylation profiles at diagnosis (Fig. 4c and Supplementary Table 7). Specifically, mmSWpos samples showed higher methylation in regulatory regions critical for transcriptional control, including TSS1500, TSS200, and first exon (Fig. 4d), suggesting a larger proportion of epigenetically repressed genes compared with mmSWneg. When we examined these regions in relation to CpG island (CGI) context, we observed a different distribution of hypermethylated probes between groups (Fig. 4e). mmSWpos samples showed increased methylation within canonical CGIs, consistent with direct gene silencing. Remarkably, Gene Set Enrichment Analysis (GSEA) of hypermethylated promoter-associated probes revealed a significant enrichment for genes targeted by the Polycomb Repressing Complex in mmSWpos samples (Fig. 4f and Supplementary Table 8-9). This observation mirrors the transcriptional regulatory patterns identified by pySCENIC analysis at Dx and D15, where multiple regulons specifically repressed at Dx and activated at D15 were directly modulated by PRC2-associated factors, including *IRX4*, *NKX2-8*, *IRX5*, *HOXD8*, *MSC*, *ZNF503*, *EGR2*, *EN1*, *OLIG1*, *VDR* (Fig. 2e and Extended Data Fig.5c-d). In conclusion, the distinct methylation landscape at Dx in mmSWpos BCP-ALL supports the peculiar immunophenotypic and transcriptomic plasticity of this subgroup. Altogether, these data suggest a central role for DNA methylation and epigenomic remodeling in the regulation of the mmSW phenomenon.

## DISCUSSION

Transient mmSW is an intriguing immunophenotypic phenomenon observed in specific genetic BCP-ALL subtypes. Despite the immunophenotypic changes detected by MFC and the treatment recommendations already provided^10^, no clear information on the biological bases are available to date. Understanding the biology of mmSW may have critical implications for patient management. *DUX4*-rearranged pediatric BCP-ALL, a molecular ALL subtype more prone to mmSW than others, usually shows a suboptimal response to induction therapy with consequent allocation in the high-risk therapeutic group, but a final favorable outcome after MRD-modulated treatment^10^. To gain insights on the biological bases of transient mmSW in BCP-ALL we applied a multiomic single-cell approach. We showed that mmSW reflects lineage plasticity within a transcriptionally unstable leukemic cell compartment that is already present at diagnosis in mmSWpos patients. Single-cell projection onto healthy hematopoietic reference further revealed the heterogeneous distribution of mmSWpos blasts along the hematopoietic developmental trajectory. Consistent with our findings, it has been recently demonstrated that a subset of BCP-ALLs, specifically *DUX4*-r and *ZNF384*-r subtypes, harbors transcriptionally unstable leukemic compartments with multipotent lineage potential and heterogeneous positioning along hematopoietic developmental trajectories, supporting the plasticity within these subtypes.^13^ Despite the predominance of *DUX4*-r subtype in our cohort, blasts at diagnosis exhibited marked inter-patient heterogeneity at the transcriptomic level, but clustered according to mmSW status. Even within the *DUX4*-r subgroup, mmSWpos and mmSWneg samples formed distinct clusters, indicating that mmSW is not a direct consequence of the underlying genetic lesion, but rather of another layer of regulation. This is also in line with the previously described MFC-MRD behavior. Although the *DUX4-r* subgroup shows a high propensity for the lineage switch, this is not unequivocal in all cases^10^.

The longitudinal analysis of matched Dx-D15 samples provided mechanistic insights into how blast cells evolved during induction therapy. In mmSWpos patients, D15 blasts loose B-cell identity and acquire a monocyte-like transcriptional profile, as defined by cell annotation and projection. The progressive loss of the leukemic *DUX4*-r-related signature from Dx to D15 supports the idea of a true transdifferentiation event, rather than a clonal replacement. By contrast, in the mmSWneg patient analyzed longitudinally, D15 blasts retain a B-cell progenitor identity at both the immunophenotypic and transcriptomic levels.

Our trajectory and cell dynamics analyses further refine the model of mmSW phenomenon, suggesting that transcriptional plasticity at diagnosis underlies the subsequent lineage switch. Of note, differently from immunophenotype, Dx blasts in mmSWpos patients appear to comprise at least two transcriptional states: a B-lineage–committed population and a second, more heterogeneous population enriched for early lymphoid and myeloid regulators. We define this latter population as characterized by a ‘fate-uncertain’ state, with high entropy and low cell-state density, consistent with increased developmental plasticity. Sitting at the root of differentiation branches and given its propensity to progressively acquiring myeloid features, this subpopulation can represent the cellular origin of transient mmSW, ultimately connecting Dx to D15 states. Notably, these findings indicate that mmSW arises from a pre-existing, transcriptionally unstable leukemic cell subset that is competent to adopt alternative lineage fates under the pressure of induction therapy. Rare BCP-ALL cases (not included in this cohort) showed the presence of a double population similar to mmSW cases detected by MFC already at diagnosis. Those patients received steroid treatment a few days before (data not shown), suggesting its role in mmSW phenomenon. Further, the upregulation of genes (i.e., *TGFB1* and *PLAGL1*), previously described to be associated with myeloid potential in *DUX4*-r blast cells,^14^ was restricted to a subpopulation of mmSWpos blasts at diagnosis in our cohort. Notably, those cells also showed the downregulation of *DUX4*-r signature, suggesting that the plasticity of *DUX4*-r blasts might not be completely guided by *DUX4* expression.

The dynamics of this process appear heterogeneous across patients but converge on a shared mechanism, suggesting that the size and timing of expansion of this plastic population at Dx can influence mmSW detection. At the regulatory level, pySCENIC analysis highlights a coordinated remodeling of gene regulatory networks during the switch. mmSWpos samples share D15-specific regulons driven by TFs implicated in HSC biology and AML pathogenesis. These regulons are relatively inactive at diagnosis but were found to be active at D15 of treatment, concomitant with myeloid-lineage reprogramming, suggesting that induction therapy unmasks or reinforces myeloid-primed regulatory programs in the plastic leukemic cell population. In parallel, regulons enriched for targets of PRC components are also activated at D15, consistent with an epigenetically controlled reconfiguration of lineage programs. Supporting the myeloid potential, a subgroup of *DUX4*-r ALLs has been shown to exhibit a chromatin accessibility profile similar to monocyte cells^14^. This is further reinforced by the presence of Ras pathway mutations in mmSWpos patients at diagnosis, a common feature of myeloid and early T-cell precursor (ETP) leukemia^21^.

Together, these findings support a model in which mmSWpos BCP-ALLs are intrinsically prone to lineage plasticity, being shaped by specific mutational and epigenetic backgrounds. Transformation of initial immunophenotype can vary based on the acquisition and/or loss of lineage defining antigens. For careful assessment of immunophenotypic changes, Project EVOLVE has recently introduced the term “lineage drift” (LD) referring to gradual immunophenotypic evolution of blast without sufficient immunophenotypic changes to meet LS definition criteria^5^.

Here we describe multilineage plasticity with mmSW during induction, related not only, as previously described to specific genetic subtypes, but also to transcriptional regulation partially defined at diagnosis and partially induced by epigenetic changes under treatment.

In conclusion, our study supports a model in which transient mmSW reflects lineage plasticity of a pre-existing, immature, fate-uncertain leukemic compartment, with implications for disease classification, response to therapy monitoring, and potential escape to B-lineage immuno-targeted therapies. Future studies should address the possible role of the immune microenvironment on phenotypic changes and under the selective pressure of antigen-directed immunotherapy (CAR-T cells and bispecific antibodies) that is revolutionizing ALL treatment landscape. With the excellent results of CD19-targeted therapy in BCP-ALL^22,23^ and the increasing use also in frontline trials, the antigen escape and lineage switch are becoming of particular interest to quickly identify patients no longer addressable by antigen-directed immunotherapy. Our data may constitute a starting point for improving the biological bases of other subtypes of lineage switch in leukemias, as well as building *in vitro/in vivo* models for targeted-therapy studies.

## MATERIALS AND METHODS

### Human samples

Bone marrow (BM) samples included in the study were collected at diagnosis (n = 20) and at Day+15 (D15) of induction therapy (n = 6) from BCP-ALL pediatric patients enrolled in AIEOP-BFM ALL frontline clinical trials and following registry phases. BCP-ALL diagnosis was based on morphology, immunophenotyping, and genetic analyses^2,11,12,24^. According to the treatment protocols, response to therapy was evaluated by Multiparametric Flow Cytometry (MFC) in BM samples collected at D15 (MFC-MRD). BM specimens were processed with LymphoSep™ (Biowest, Nuaillé, France) gradient centrifugation to isolate mononuclear cells from which DNA and RNA were extracted for genetic investigations. Written informed consent to use residual diagnostic material for research purposes was obtained from parents or legal guardians, in compliance with the Declaration of Helsinki and local institutional ethical committee. Patients enrolled in the study were selected considering the availability of residual material at diagnosis and at D15. Initial sample selection was guided by mmSW at D15, resulting in an overrepresentation of *DUX4*-rearranged cases in our cohort.

### Multiparametric flow cytometry

BM samples collected at diagnosis and at D15 were analyzed according to previously established standard operating procedures^11,24^. Immunophenotyping was performed on erythrocyte-lysed whole BM samples. Antigen expression was graded as negative, weak-positive, strong-positive, or partial-positive by comparing the fluorescence intensity and blast distribution pattern to a negative control, following the AIEOP-BFM 2016 Consensus Guidelines for ALL immunophenotyping^12^. BCP-ALL was defined by the presence of a blast population strongly positive for at least two antigens among CD19, CD10, intracellular CD22 (iCD22), and intracellular CD79a (iCD79a). Patients were classified according to the European Group for the Immunological Characterization of Leukemias (EGIL) classification (B-I, B-II, B-III, and B-IV)^12,25^. CD371 expression was evaluated using CD371-PE (clone 50C1, Becton Dickinson, Franklin Lakes, NJ) for diagnostic immunophenotyping, and CD371-PC5.5 (clone 50C1, BioLegend, San Diego, CA) included in the dry 10-color preformulated DuraClone 10 Color Custom Mix (Beckman Coulter, Inc, Brea, CA) for MFC-MRD monitoring^10^. MFC-MRD assessment was performed at D15 according to the therapeutic protocols, and samples with a cluster of at least 10 events showing leukemia-associated immunophenotypic features were defined as MRD positive^11,12^. In addition, we evaluated the presence of a myelomonocytic switch at D15 as previously described^10^.

### Single-cell library preparation and sequencing

A multiomic single-cell approach was applied to BM samples collected at diagnosis (n = 11) and at D15 (n = 6) from pediatric patients with BCP-ALL. All samples utilized were leftover from routine diagnostic and MRD procedures. Frozen BM samples were thawed, assessed for cell viability, and counted using the Trypan Blue exclusion test (Sigma-Aldrich, St. Louis, MO, USA). One million cells per sample with a cell viability > 65% were stained for 30 min on ice with Sample Tags (ST; BD Biosciences, San Jose, CA, USA) and AbSeq oligonucleotide-conjugated antibodies (AbSeq; BD Biosciences, San Jose, CA, USA), protected from light. We designed a panel of 14 AbSeq antibodies targeting the same surface markers used in routine diagnostic immunophenotyping and MFC-MRD in BCP-ALL as defined by the AIEOP-BFM 2016 Consensus Guidelines for ALL immunophenotyping^12^. In detail, we selected: CD19 (SJ25-C1), CD20 (L27), CD10 (HI10A), CD58 (1C3), CD33 (P67.6), CD34 (8G12), CD38 (HB-7), CD371 (50C1), CD2 (RPA-2.10), CD3 (SK7), CD11b (ICRF44), CD14 (MΦP-9), CD15 (W6D3), and CD45 (HI30) AbSeq. After staining, single cells were captured using BD Rhapsody (BD Biosciences, San Jose, CA, USA) single-lane cartridges according to the manufacturer’s instructions. For each experiment, 3 libraries (ST, AbSeq, and whole transcriptome [WTA]) were generated and sequenced on a NovaSeq 6000 platform (Illumina, San Diego, CA, USA) using S2 or S4 flow cell (paired-end, 2×100 cycles; 75 cycles for read 1 and 125 cycles for read 2).

### Single-cell multiomics primary and secondary data analyses

FASTQ files generated from multiomic single-cell sequencing experiments were processed using the BD Rhapsody Sequencing Analysis pipeline (Velsera, https://www.sevenbridges.com) using default parameters. The pipeline uses Bowtie2 to map filtered read 2 (R2) sequences to the human reference genome (GRCh38). Reads that align to a Sample Tag sequence and are associated with a putative cell are used to assign each cell to its corresponding sample. The resulting DBEC-corrected unique molecular identifier (UMI) count matrices were used for downstream analyses. Quality control and pre-processing were performed using Seurat (version 5.4.0)^16^. We applied filtering with the PercentageFeatureSet() function to retain cells with: (i) a low percentage of reads mapping to mitochondrial genes (< 39%, run-dependent) and (ii) number of detected mRNA molecules between 500 and 5,000 per cell. Data were normalized using the global-scaling method LogNormalize and scaled and linearly transformed using the ScaleData() function. The resulting corrected counts were used for visualization and clustering analyses. The analysis workflow, including specific parameters, was tailored to each sequencing run and group of samples to account for technical variability. Principal component analysis (PCA) was first performed on the processed expression matrices. Cell clustering was then carried out by constructing a k-nearest neighbors (KNN) graph based on Euclidean distance in PCA space and refining edge weights between cell pairs according to the overlap of their local neighborhoods (Jaccard similarity). Uniform Manifold Approximation and Projection (UMAP) was subsequently used for dimensionality reduction and visualization. UMAPs were plotted using a customized function of the monocle3 (version 1.4.13)^26^ R package. Density UMAPs for AbSeq intensities were generated using the Nebulosa package (version 1.0.1)^27^. Pseudo-bulk count matrices were generated from single-cell data using the Seurat function AggregateExpression(), iterating over each sample’s gene expression matrix. All analyses, unless otherwise specified and comprising the ones mentioned below, were performed using R (version 4.2.2) on Ubuntu 22.04 using an Intel vPro i9 processor. Utilities R packages were used to preprocess and tidy up data (i.e. tidyverse [version 2.0.0]^28^, data.table [version 1.18.2.1]^29^, patchwork [version 1.2.0]^30^, ggtext [version 0.1.2]^31^, ggpubr [version 0.6.0]^32^ and symphony [version 0.1.1]^33^).

### Cell type annotations and non-malignant population

Single cells were annotated using Azimuth BM reference^16,34,35^. Markers for each non-malignant cluster were calculated using FindMarkers() in Seurat (version 5.4.0)^16^ using min.pct = 0.5. A full list of markers for each cluster is reported in Supplementary Table 10. The top 10 up-regulated genes for each cluster (average log2 fold-change) were used to calculate non-malignant scores with the AddModuleScore() function (Seurat package).

### CNVs inference on single-cell transcriptome

Copy number variations (CNVs) were inferred from filtered single-cell gene expression count matrices using a containerized version of inferCNV (of the Trinity CTAT Project, https://github.com/broadinstitute/inferCNV, version 1.21.0) on Singularity (apptainer, version 1.3.1) in an HPC cluster. Seurat clusters were used as annotations, labeling also previously identified non-malignant cells clusters, used as reference ‘normal’ cells. A cutoff of 0.1 was set for the infercnv::run function, with denoise and HMM set to TRUE.

### Identification of blast developmental stages

Projection of single cells onto a reference atlas of healthy human hematopoiesis was performed using the BoneMarrowMap package (https://github.com/andygxzeng/BoneMarrowMap) in R. Cells were first projected onto the global reference map comprising 263,159 hematopoietic cells spanning from stem and progenitor compartments through all differentiated BM lineages^15^. Subsequently, B-cell lineage cells were subset and re-projected onto a B-cell developmental reference atlas using customized scripts derived from the BoneMarrowMap package^13^.

### Transcriptional priming and *DUX4*-r signature score

The AddModuleScore() function (Seurat package) was applied on single-cell WTA data to calculate transcriptional priming towards stemness^16^ and myeloid lineage^17^ for each single cell. The density plot was generated using ggplot2 (version 3.5.0)^36^. The same approach was used to calculate the *DUX4*-r signature score based on gene expression profile (only the up-regulated genes) of this BCP-ALL subgroup according to previously identified signatures, used in the ALLCatchR package (https://github.com/ThomasBeder/ALLCatchR)^18^.

### pySCENIC regulatory network

Regulatory network inference and transcription factor activity analysis were performed using a containerized version of pySCENIC^37^ (version 0.12.1) in Singularity (apptainer, version 1.3.1) in an HPC cluster. Gene regulatory networks (GRNs) were inferred from the single-cell expression matrices using the GRNBoost2 workflow, followed by motif enrichment analysis to define regulons. Regulon activity scores (AUCell) were then computed per cell to quantify transcription factor–target module activity across clusters and conditions. Downstream visualization and comparison of regulon activities were integrated with the Seurat-derived clustering and represented as heatmaps and UMAP embeddings.

### Pseudotime and cell-state dynamics

Force-directed layouts and pseudotime trajectories were computed from preprocessed single-cell data using Palantir (version 1.3.6)^38^ and scanpy (version 1.10.4)^39^. Trajectory start and end points were manually curated based on cell labels and marker expression patterns. Cell-state densities were inferred with Mellon (version 1.4.3)^20^, leveraging the Palantir-derived force-directed layouts. MAGIC algorithm^40^ was applied to denoise transcriptome data and obtain gene expression trends. All packages were run in Python (version 3.10.12)^41^.

### Whole Transcriptome Sequencing

Whole Transcriptome Sequencing (RNA-seq) analysis was performed for 19 BCP-ALL patients (n = 14 mmSWpos, n = 5 mmSWneg). Total RNA was extracted at diagnosis from BM mononuclear cells using the TRIzol method (ThermoFisher Scientific, Waltham, MA, USA) and libraries were prepared with the Universal Plus mRNA-Seq kit with NuQuant (Tecan, Switzerland). Sequencing was carried out on an Illumina NovaSeq X Plus platform (Illumina, San Diego, CA, USA). Multiple analysis tools were applied to the bulk RNA-seq data. The nf-core/rnaseq pipeline (version 3.18.0)^42^ in Nextflow (version 24.04.2)^43^ was used for alignment and quantification on a HPC cluster. AllCatchR (version 1.0; R version 4.5.0) was used to predict the BCP-ALL molecular subtype^18^. A containerized version of FusionCatcher (version 1.33)^44^ in the Nextflow-implemented (version 23.10.1)^43^ nf-core/rnafusion pipeline (version 3.0.2)^42^ was applied to identify somatic fusion genes. Variants were called using the nf-core/rnavar pipeline (version 1.1.0)^42^ implemented in Nextflow (version 24.04.2) on an HPC cluster and annotated with VEP (version 111.0)^45^. Variants that were flagged as PASS were retained and further filtered by requiring an AD > 3 and DP > 10. These variants were then filtered for high or moderate predicted functional impact, as annotated by SnpEff (version 5.1d)^46^, and for a gnomAD^47^ allele frequency < 0.001. Additional filtering included only those affecting genes in a curated list of 376 genes frequently altered in BCP-ALL^48^. Genes without an assigned pathway in the original list were manually annotated. Pathways assignment has been adapted by collapsing and renaming similar networks (Supplementary Table 11). Variants were manually reviewed by inspecting BAM files.

### Whole Genome Sequencing

Genomic DNA was extracted and purified by using the Qiagen (Hilden, Germany) DNA extraction kit according to the manufacturer’s instructions from BM mononuclear cells (n = 13 mmSWpos, n = 6 mmSWneg BCP-ALLs). For sequencing library preparation TruSeq Illumina DNA Nano kit was used and then sequenced by Illumina sequencer (Illumina, San Diego, CA, USA). The nf-core/sarek pipeline (version 3.4.4)^42^ in Nextflow (version 24.04.2)^43^ on an HPC cluster was applied to process FASTQ files and reads were mapped onto the human reference (GRCh38). Variant calling was performed using Mutect2 (GATK version 4.5.0.0)^49^ and only PASS variants were kept, while the annotation was done by VEP (version 111.0)^45^ and SnpEff (version 5.1d)^46^. Only variants with a high and moderate predicted impact according to SnpEff and with a gnomAD^47^ frequency < 0.001 were considered. Next, as per RNA-seq data, the list of 376 recurrently altered genes in BCP-ALLs was used to filter variants^48^. Genes without an assigned pathway in the original list were manually annotated. Pathways assignment has been adapted by collapsing and renaming similar networks (Supplementary Table 11). The R programming language (version 4.5.0)^50^ was used for downstream data analysis and visualization. The input variants files were imported using readxl (version 1.4.5)^51^. After integration of variants from both WGS and RNAseq datasets, the oncoplot was generated using the oncoPrint function from the ComplexHeatmap package (version 2.24.1)^52^, with graphical support from circlize (version 0.4.17)^53^. The heatmap was generated using pheatmap (version 1.0.13)^54^. Data preprocessing and manipulation were performed using tidyverse (version 2.0.0)^28^, dplyr (version 1.2.0)^55^ and tibble (version 3.3.1)^28^ packages. Additional graphical customization and annotations were implemented using grid (version 4.5.0)^56^, gtable (version 0.3.6)^57^, RColorBrewer (version 1.1-3)^58^, showtext (version 0.9-7)^59^ and magick (version 2.9.0)^60^.

### DNA methylation profiling

Genomic DNA extracted from BM samples at diagnosis from our cohort of 19 (n = 14 mmSWpos, n = 5 mmSWneg) BCP-ALL pediatric patients was processed for methylation analysis. We adopted Illumina Infinium MethylationEPIC BeadChip system (bisulfite-based amplicon arrays, Illumina, San Diego, CA, USA) according to the manufacturer’s protocol to assess DNA methylation at genome-wide level. Raw methylation data (IDAT files) were obtained by the iScan System (Illumina, San Diego, CA, USA) and then processed by the CHAMP package (version 2.38.0)^61^ in R (version 4.2.2)^50^. At first, the quality check step filtered out probes with detection P value > 0.01, probes with less than 3 beads in at least 5% of samples per probe, probes related to SNPs, and probes located in X and Y chromosomes. After filtering we obtained a beta matrix, where the beta values are defined as continuous variables between 0 and 1, representing the ratio of the intensity of the methylated bead type to the combined locus intensity. We applied a BMIQ (Beta MIxture Quantile dilation) normalization and Combat batch correction for Slides parameter. The top 1,000 variable probes were used to generate the multidimensional scaling (MDS) plot. Then, Differentially Methylated Probes (DMPs) were calculated using a linear model (as implementation of limma package) by comparing mmSWneg to mmSWpos samples. DMPs having an absolute delta beta value > 0.3 were selected for further analyses. Benjamini-Hochberg method was used and significant DMPs were defined with P value = 0.05. Gene Set Enrichment Analysis (GSEA) was performed using the CHAMP package, on DMPs mapping on CGIs (CpG islands) of TSS (Transcription Starting Site). Fisher’s exact test was used, setting adjusted P value = 0.05.

## DATA AND CODE AVAILABILITY

Single-cell multiomic data and methylation data are deposited on GEO under the accession GSE330258. RNA-seq data are deposited on ArrayExpress under the accession E-MTAB-17083. Custom scripts and parameter settings are available at: https://github.com/albertopeloso/BCPALL-MMSwitch-SingleCellLandscape.

## AUTHOR CONTRIBUTIONS

G.G. contributed to study design, performed experiments, analyzed data, generated figures and conceptualized and wrote the manuscript; A.P. contributed to study design, implemented bioinformatic pipelines, analyzed data, generated figures and conceptualized and wrote the manuscript; E.V. conceived and designed the study, contributed to sample collection, provided clinical data and wrote the manuscript; P.S. performed experiments and analyzed data; M.V. analyzed data and generated figures; C.F. performed experiments and analyzed data; A.C. performed experiments; G.F. and G.C. contributed to sample collection and provided biological and molecular clinical data; F.L., C.R., F.M., A.M.T. and A.B. provided samples and clinical data; M.N.D. provided expertise and critical discussion; S.B. conceived, designed and supervised the study, conceptualized and wrote the manuscript; B.B. conceived, designed and supervised the study, conceptualized and wrote the manuscript and provided funding. All authors reviewed and approved the manuscript.

## ACKNOWLEDGMENTS

We would like to thank the members of the Division of Pediatric Hematology, Oncology and Stem Cell Transplant, Women’s and Children’s Health Department for the diagnostic activity. The authors would like to thank all the patients involved in the study, their families, and the clinical teams who took care of them. This work is in memory of Prof. Giuseppe Basso (1948–2021).

