## Extended Data Figures for "Single-cell multiomic profiling reveals lineage plasticity in pediatric B-lineage Acute Lymphoblastic Leukemia during the early phase of treatment"

##### Extended Data Fig. 1: Selection of BCP-ALL blasts at diagnosis by multi-omic data integration

**a-b,** UMAPs showing 35,127 cells from 11 BCP-ALL BM samples (n = 3 mmSWneg and n = 8 mmSWpos) at diagnosis. In (**a**) colors represent the different patients. In (**b**) cells are colored by annotation. Annotated cells were colored based on their ontogeny: dendritic cells (ASDC, cDC2, pDC, pro-mDC, pre-pDC) in magenta; Erythroid cells (EMP, Early Eryth, Late Eryth, Prog Mk) in orange; HSC in red; Mature B-cells (Memory B, Naive B, Plasma) in blue; Monocytes (CD14 Mono, CD16 Mono) in lilac; B-cell precursors (CLP, pre B, pro B, transitional B) in light blue; Precursor myeloid cells (BaEoMA, GMP) in purple; Stromal cells in yellow; T-cells (CD4 Memory, CD4 Naive, CD8 Effector_1, CD8 Effector_2, CD8 Memory, MAIT) in green. **c,** Co-expression of canonical B-cells, myeloid cells and T-cells surface markers, used to identify non-malignant populations. **d,** Dotplot showing top 10 differentially expressed genes among non-malignant cells. Dot size reflects the percentage of cells expressing each gene and colour indicated average log2 fold-change (log2FC) between each non-malignant cell cluster and all the other cells in the dataset. **e,** Gene-set scores for B-cell, myelo/erytrhroid and T-cell programs. **f,** Single-cell copy-number profiles inferred by inferCNV from each patient, ordered by genomic position, revealing patterns of copy-number gains (red) and losses (blue). Rows represent non-malignant and leukemic cells, the latter divided by Seurat clustering.

##### Extended Data Fig. 2: Characterization of BCP-ALL blasts at diagnosis by AbSeq surface markers

Transcriptomic-guided 2D UMAP representation of the expression of 13 surface markers (AbSeq) on blast cells from patients at diagnosis. Density is represented as color gradient. CD3 AbSeq was profiled only in BCPALL#11.

##### Extended Data Fig. 3: Projection onto healthy BM reference atlases

**a-b,** Dimensionality reduction maps of the two healthy reference datasets used for projection and labeling. In reference (**a**) all BM cell populations are represented[15], while (**b**) dataset is specifically focused on B-cell lineage differentiation[13]. **c-d,** Projection of single cells from patients onto the B-cell lineage reference map. Each UMAP represents one patient, with mmSWneg patients grouped in (**c**) and mmSWpos ones in (**d**).

##### Extended Data Fig. 4: Selection of BCP-ALL blasts in coupled Dx-D15 samples by multi-omic data integration

**a-b,** UMAP of the integrated single-cell transcriptomes from 8 patients with matched Dx-D15 samples, coloured by sample of origin in (**a**) and timepoint in (**b**). **c,** Joint density of B-, myelo/erythroid and T-cells markers expression to subset non-malignant cells at the two timepoints. Joint density is plotted as shades of blue. **d,** Gene-set scores for non-malignant compartments at the two timepoints. Colors represent markers score. **e,** Expression of surface markers (AbSeq) in blasts at Dx and D15. CD3 was profile only in BCPALL#11. Colors are indicative of density.

##### Extended Data Fig. 5: Characterization of BCP-ALL blasts in coupled Dx-D15 samples by UMAP and pySCENIC

**a-b,** UMAPs of blast cells from each patient with matched Dx-D15 samples, coloured by sample and cell type. MmSWpos patients are depicted in (**a**), while the mmSWneg patient in (**b**). Colors indicate BM cell subpopulations **c,** Harmony-integrated UMAP generated from pySCENIC regulons data layer. Cells are colored by sample/ time point. **d,** pySCENIC regulon activity of selected regulons within the leukemic populations in patients at Dx and D15. Color gradient represents activity score.

##### Extended Data Fig. 6: FDL emphasizes the evolutionary trajectory in the Dx-D15 axis

**a,** Force-directed layout (FDL) embeddings of Dx and D15 leukemic blasts from four patients, showing a trajectory between the two timepoints. **b,** *DUX4*-r signature score projected onto FDL. Color gradient represent the signature score. **c-f,** MAGIC-imputed gene diffusion maps of selected genes, highlighting their dynamic evolution on the Dx-D15 axis. Color gradient indicates MAGIC-imputed gene expression score.

##### Extended Data Fig. 7: Pseudotime and trajectory between the two timepoints

**a-d,** FDLs showing inferred pseudotime and Palantir entropy on the Dx-D15 axis of four mmSWpos patients. Pseudotime ranging from 0 to 1 is depicted as color gradient on the left side of each coupled plot, while entropy values are shown as color gradient on the right side. **e,** Mellon clipped density values projected on FDL representations of previously reported patients at Dx and D15. Color gradient is indicative of density values. **f-i,** Identification of two major branches in each patient, corresponding to 'B cell' branch, connecting two populations at Dx, and 'Mono' branch ranging from Dx to the monocytoid blast population at D15. **j-m,** Gene trends maps of selected genes along pseudotime and the Dx-D15 axis in each patient. Mono and B-cell trajectories are depicted in green and orange, respectively.

##### Extended Data Fig. 8: Genomic landscape of BCP-ALL at diagnosis

Oncoplot showing variants identified in BCP-ALL patients at Dx by WGS and RNAseq (n = 6 mmSWneg and n = 15 mmSWpos). MmSWneg are reported in blue, while mmSWpos in orange. Purple and yellow indicate the source of data, being WGS and RNAseq, respectively. Variant types are reported in different colors, based on their classification. In each cell, a bold black border and round circle indicates that the variant was identified on RNAseq and WGS data, respectively.

#### Extended Data Fig. 1: Selection of BCP-ALL blasts at diagnosis by multi-omic data integration

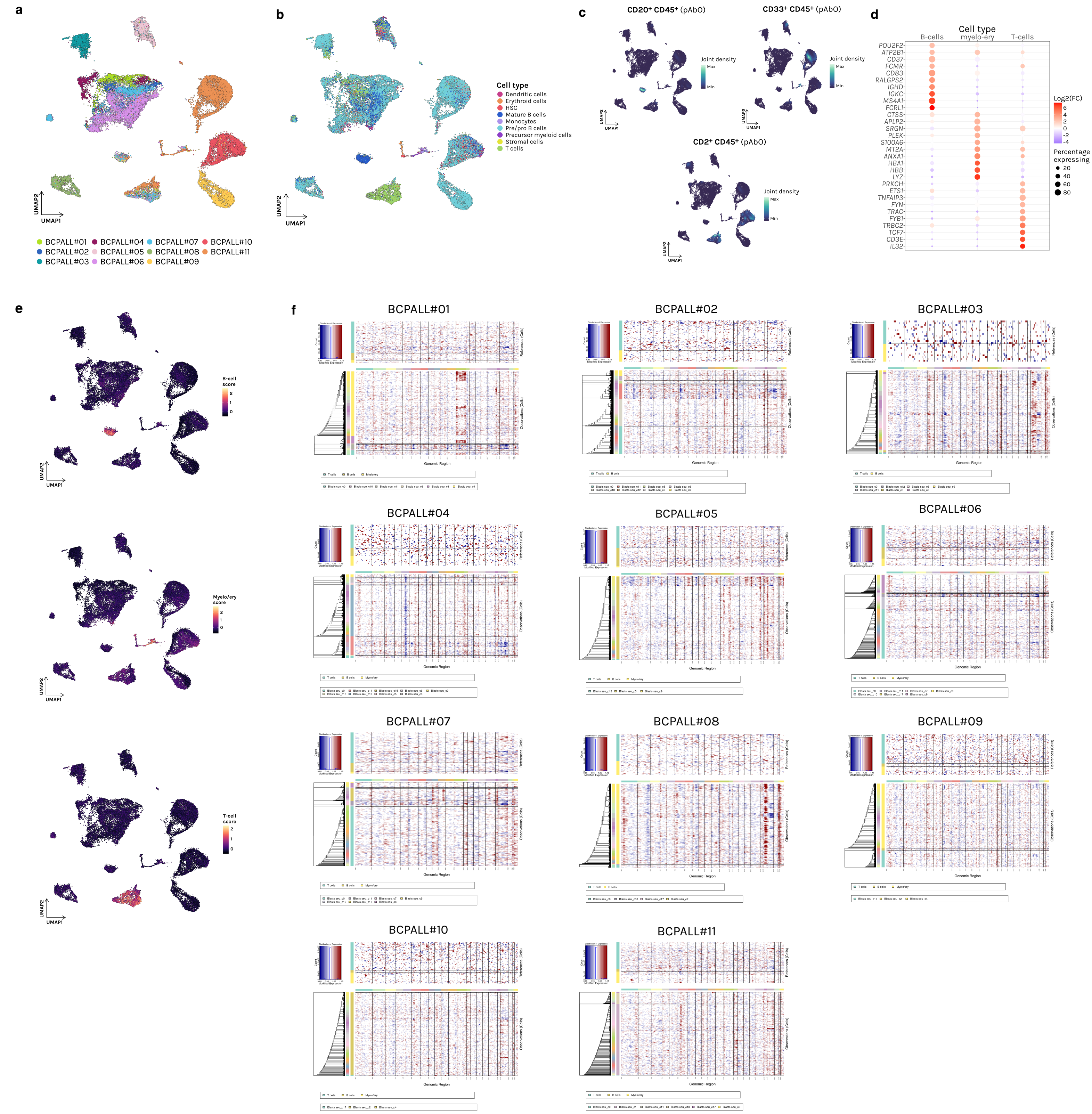


**a-b,** UMAPs showing 35,127 cells from 11 BCP-ALL BM samples (n = 3 mmSWneg and n = 8 mmSWpos) at diagnosis. In (**a**) colors represent the different patients. In (**b**) cells are colored by annotation. Annotated cells were colored based on their ontogeny: dendritic cells (ASDC, cDC2, pDC, pro-mDC, pre-pDC) in magenta; Erythroid cells (EMP, Early Eryth, Late Eryth, Prog Mk) in orange; HSC in red; Mature B-cells (Memory B, Naive B, Plasma) in blue; Monocytes (CD14 Mono, CD16 Mono) in lilac; B-cell precursors (CLP, pre B, pro B, transitional B) in light blue; Precursor myeloid cells (BaEoMA, GMP) in purple; Stromal cells in yellow; T-cells (CD4 Memory, CD4 Naive, CD8 Effector_1, CD8 Effector_2, CD8 Memory, MAIT) in green. **c,** Co-expression of canonical B-cells, myeloid cells and T-cells surface markers, used to identify non-malignant populations. **d,** Dotplot showing top 10 differentially expressed genes among non-malignant cells. Dot size reflects the percentage of cells expressing each gene and colour indicated average log2 fold-change (log2FC) between each non-malignant cell cluster and all the other cells in the dataset. **e,** Gene-set scores for B-cell, myelo/erytrhroid and T-cell programs. **f,** Single-cell copy-number profiles inferred by inferCNV from each patient, ordered by genomic position, revealing patterns of copy-number gains (red) and losses (blue). Rows represent non-malignant and leukemic cells, the latter divided by Seurat clustering.

#### Extended Data Fig. 2: Characterization of BCP-ALL blasts at diagnosis by AbSeq surface markers

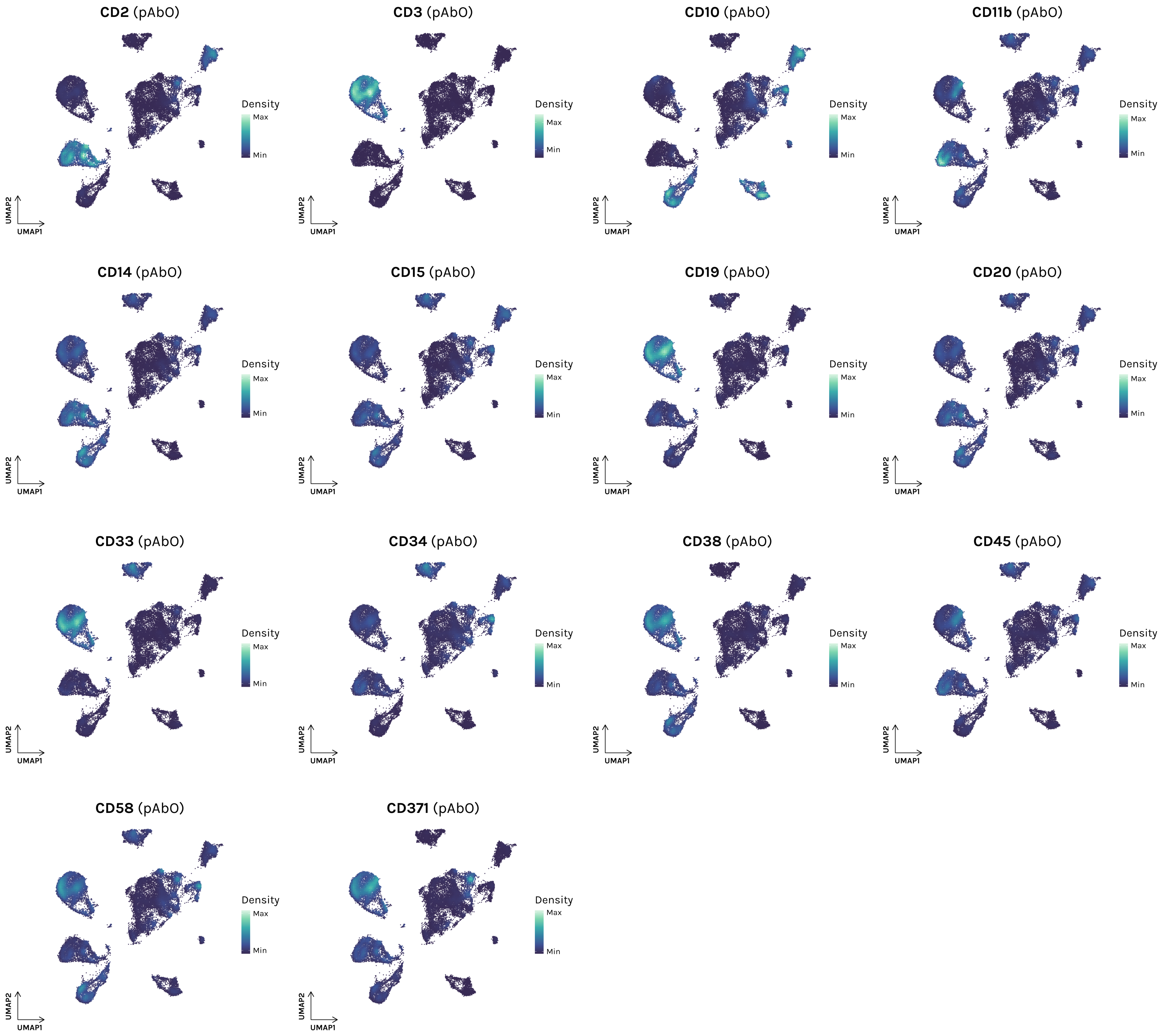


Transcriptomic-guided 2D UMAP representation of the expression of 14 surface markers (AbSeq) on blast cells from patients at diagnosis. Density is represented as color gradient. CD3 AbSeq was profiled only in BCPALL#11.

### Extended Data Fig. 3: Projection onto healthy BM reference atlases

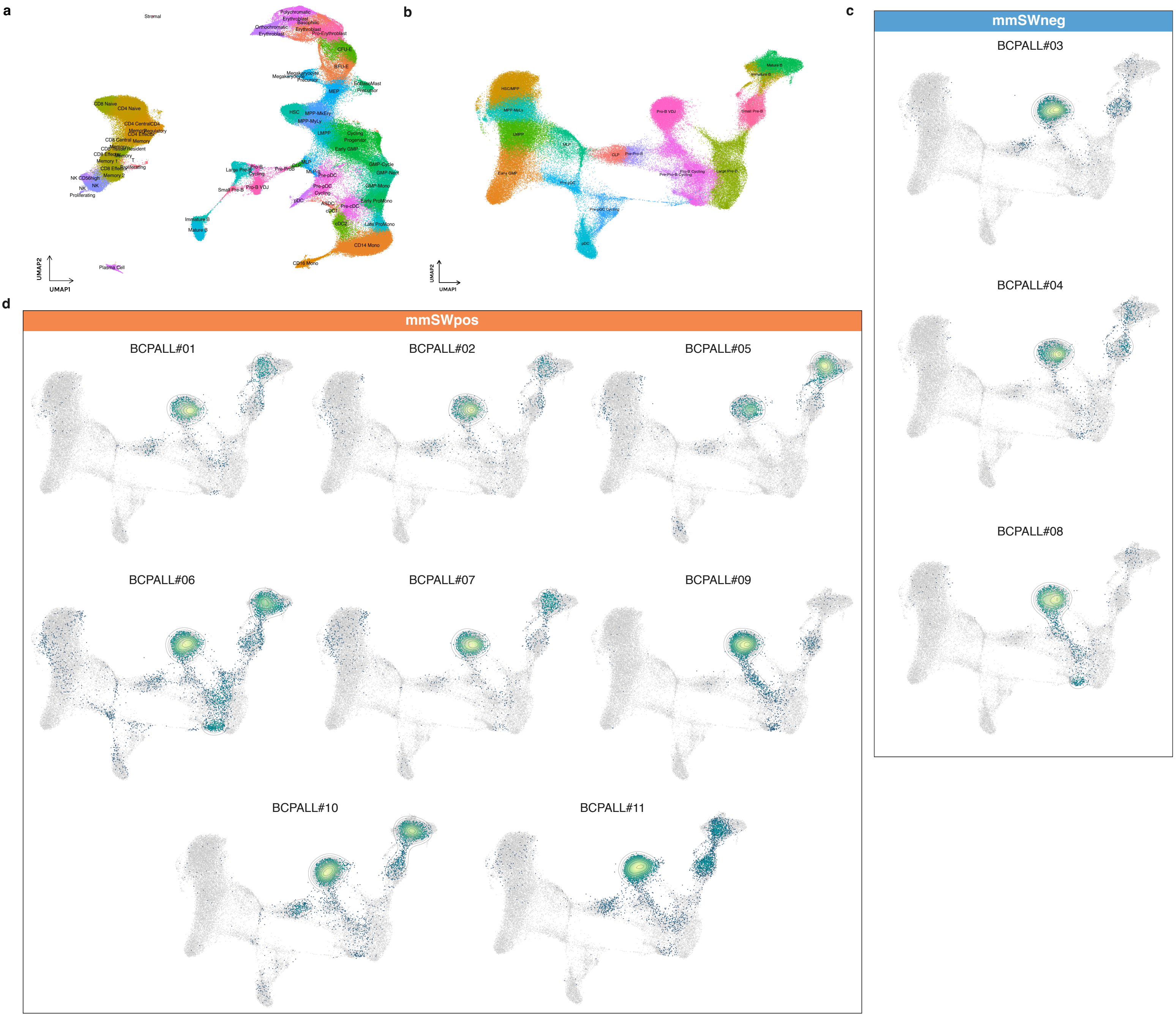


**a-b,** Dimensionality reduction maps of the two healthy reference datasets used for projection and labeling. In reference (**a**) all BM cell populations are represented[15], while (**b**) dataset is specifically focused on B-cell lineage differentiation[13]. **c-d,** Projection of single cells from patients onto the B-cell lineage reference map. Each UMAP represents one patient, with mmSWneg patients grouped in (**c**) and mmSWpos ones in (**d**).

#### Extended Data Fig. 4: Selection of BCP-ALL blasts in coupled Dx-D15 samples by multi-omic data integration

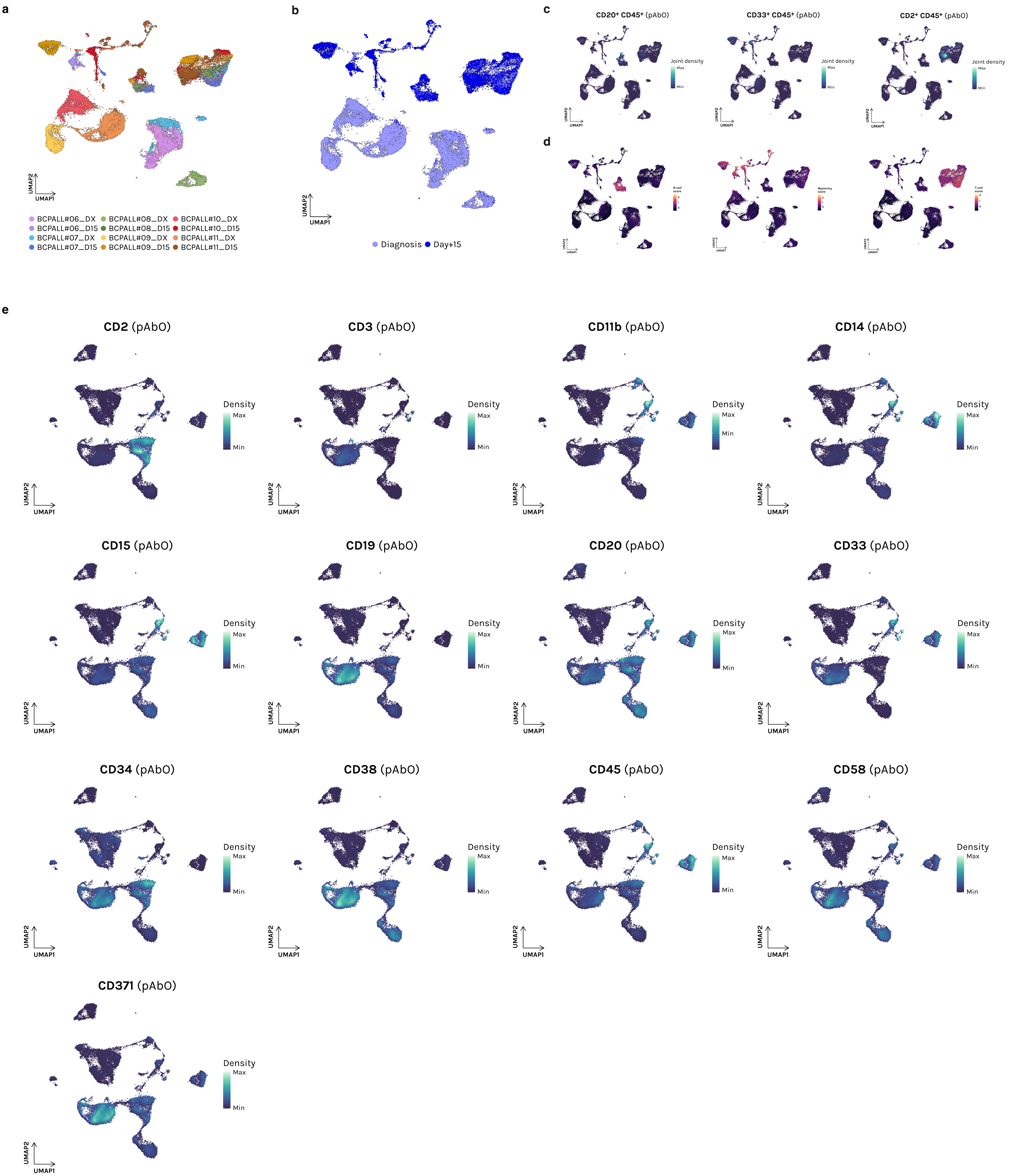


**a-b,** UMAP of the integrated single-cell transcriptomes from 8 patients with matched Dx-D15 samples, coloured by sample of origin in (**a**) and timepoint in (**b**). **c,** Joint density of B-, myelo/erythroid and T-cells markers expression to subset non-malignant cells at the two timepoints. Joint density is plotted as shades of blue. **d,** Gene-set scores for non-malignant compartments at the two timepoints. Colors represent markers score. **e,** Expression of surface markers (AbSeq) in blasts at Dx and D15. CD3 was profiled only in BCPALL#11. Colors are indicative of density.

### Extended Data Fig. 5: Characterization of BCP-ALL blasts in coupled Dx-D15 samples by UMAP and pySCENIC

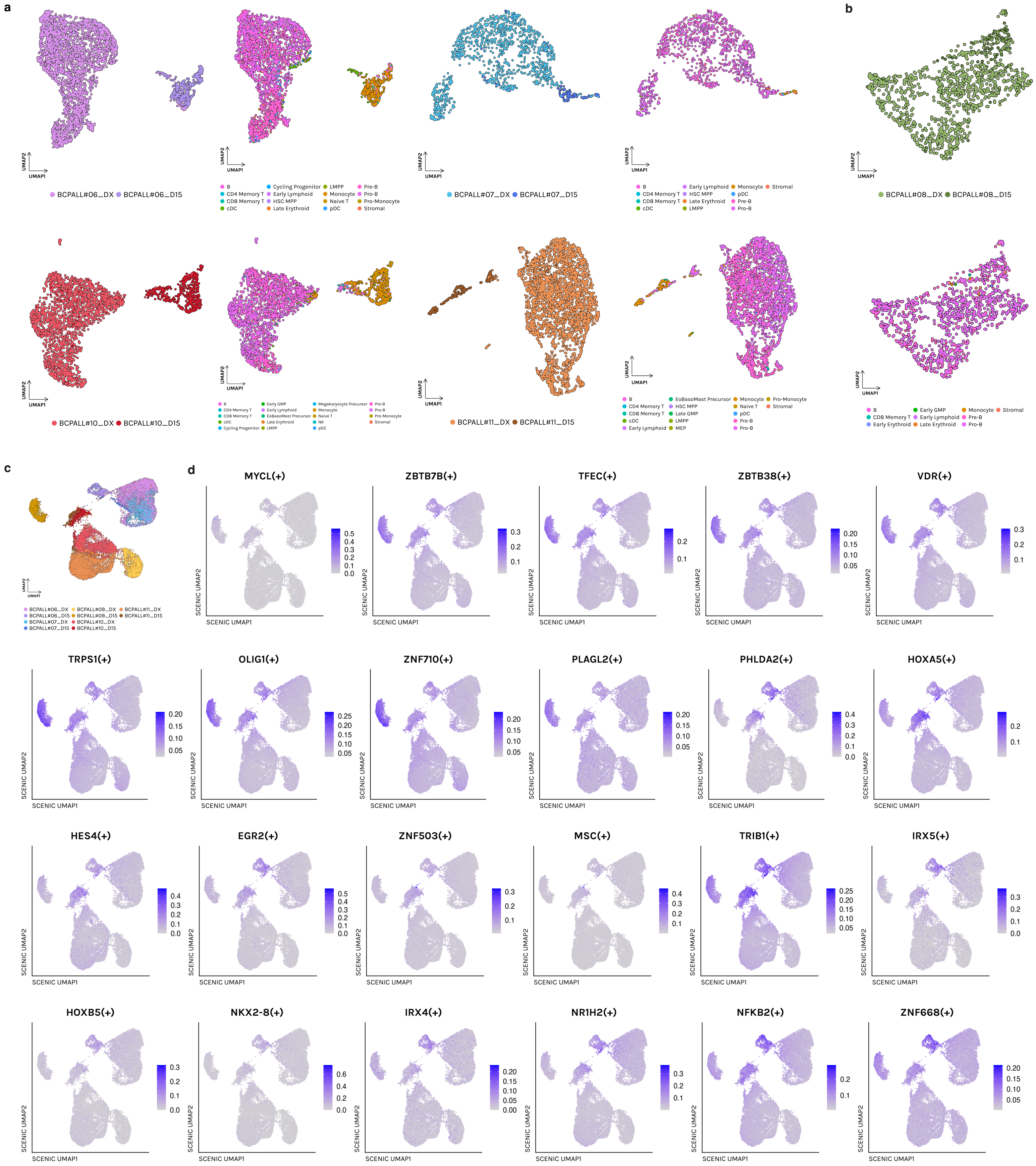


**a-b,** UMAPs of blast cells from each patient with matched Dx-D15 samples, coloured by sample and cell type. MmSWpos patients are depicted in (**a**), while the mmSWneg patient in (**b**). Colors indicate BM cell subpopulations **c,** Harmony-integrated UMAP generated from pySCENIC regulons data layer. Cells are colored by sample/ time point. **d,** pySCENIC regulon activity of selected regulons within the leukemic populations in patients at Dx and D15. Color gradient represents activity score.

#### Extended Data Fig. 6: FDL emphasizes the evolutionary trajectory in the Dx-D15 axis

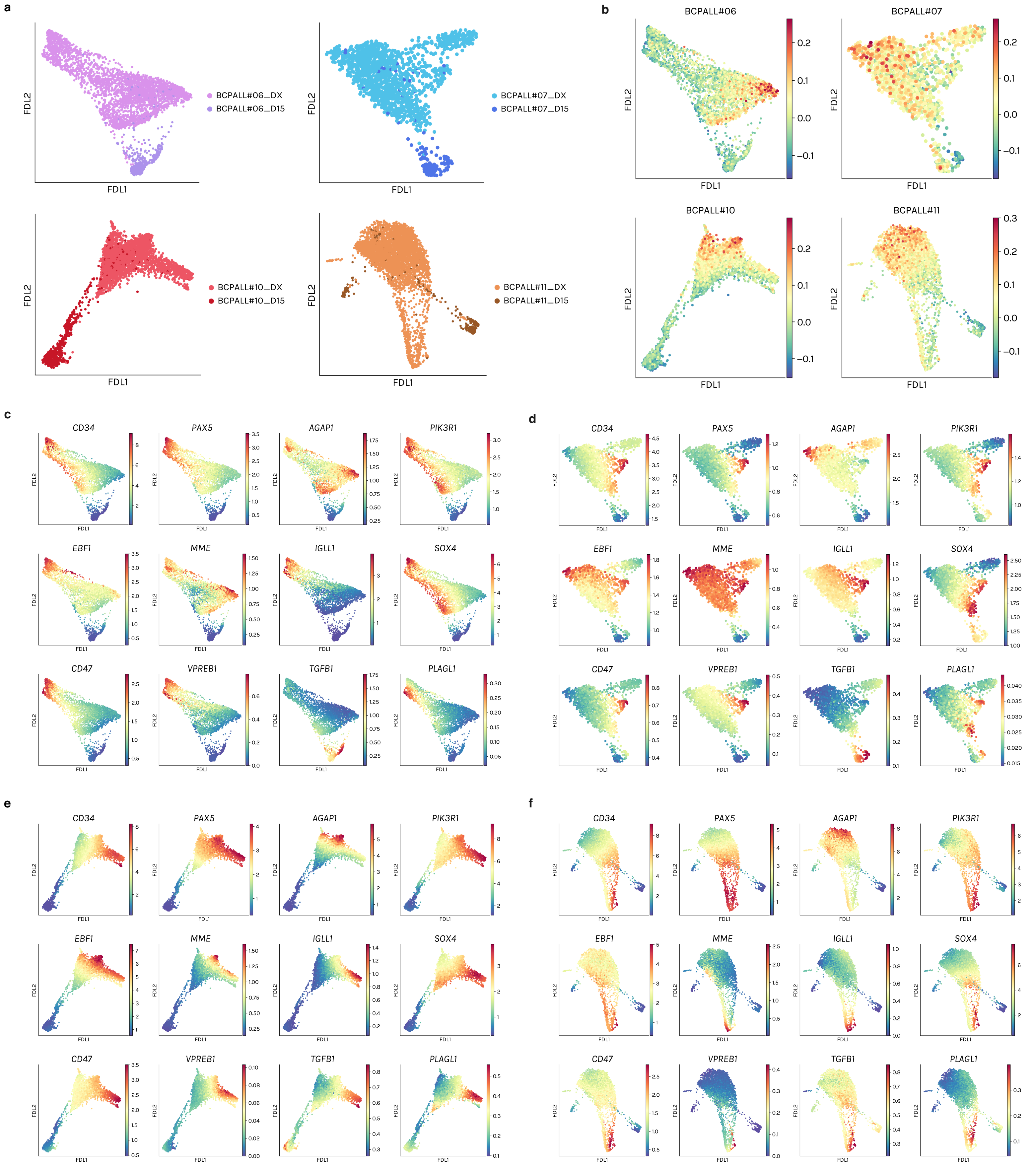


**a,** Force-directed layout (FDL) embeddings of Dx and D15 leukemic blasts from four patients, showing a trajectory between the two timepoints. **b,** *DUX4*-r signature score projected onto FDL. Color gradient represent the signature score. **c-f,** MAGIC-imputed gene diffusion maps of selected genes, highlighting their dynamic evolution on the Dx-D15 axis. Color gradient indicates MAGIC-imputed gene expression score.

### Extended Data Fig. 7: Pseudotime and trajectory between the two timepoints

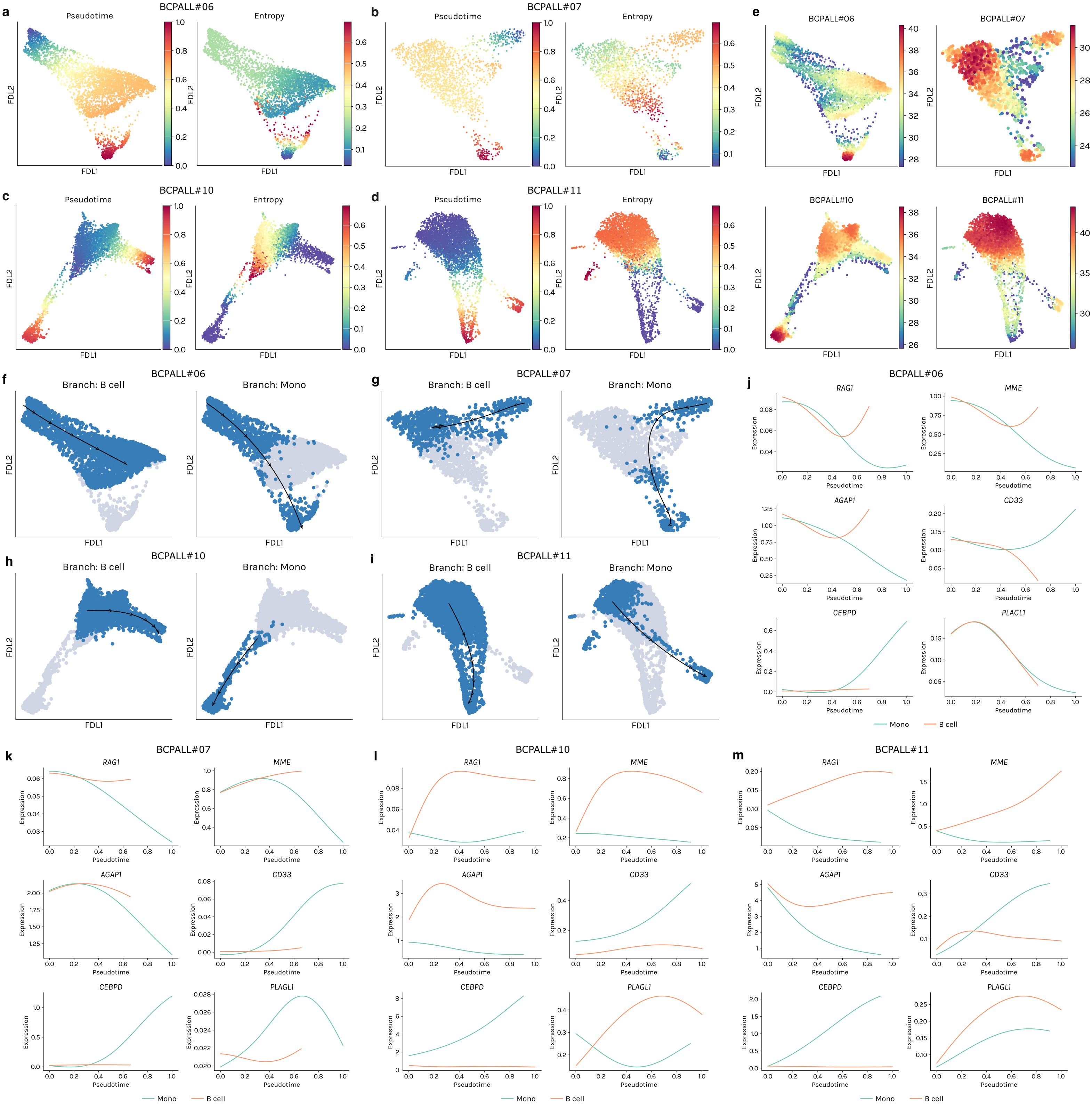


**a-d,** FDLs showing inferred pseudotime and Palantir entropy on the Dx-D15 axis of four mmSWpos patients. Pseudotime ranging from 0 to 1 is depicted as color gradient on the left side of each coupled plot, while entropy values are shown as color gradient on the right side. **e,** Mellon clipped density values projected on FDL representations of previously reported patients at Dx and D15. Color gradient is indicative of density values **f-i,** Identification of two major branches in each patient, corresponding to 'B cell' branch, connecting two populations at Dx, and 'Mono' branch ranging from Dx to the monocytoid blast population at D15. **j-m,** Gene trends maps of selected genes along pseudotime and the Dx-D15 axis in each patient. Mono and B-cell trajectories are depicted in green and orange, respectively.

#### Extended Data Fig. 8: Genomic landscape of BCP-ALL at diagnosis

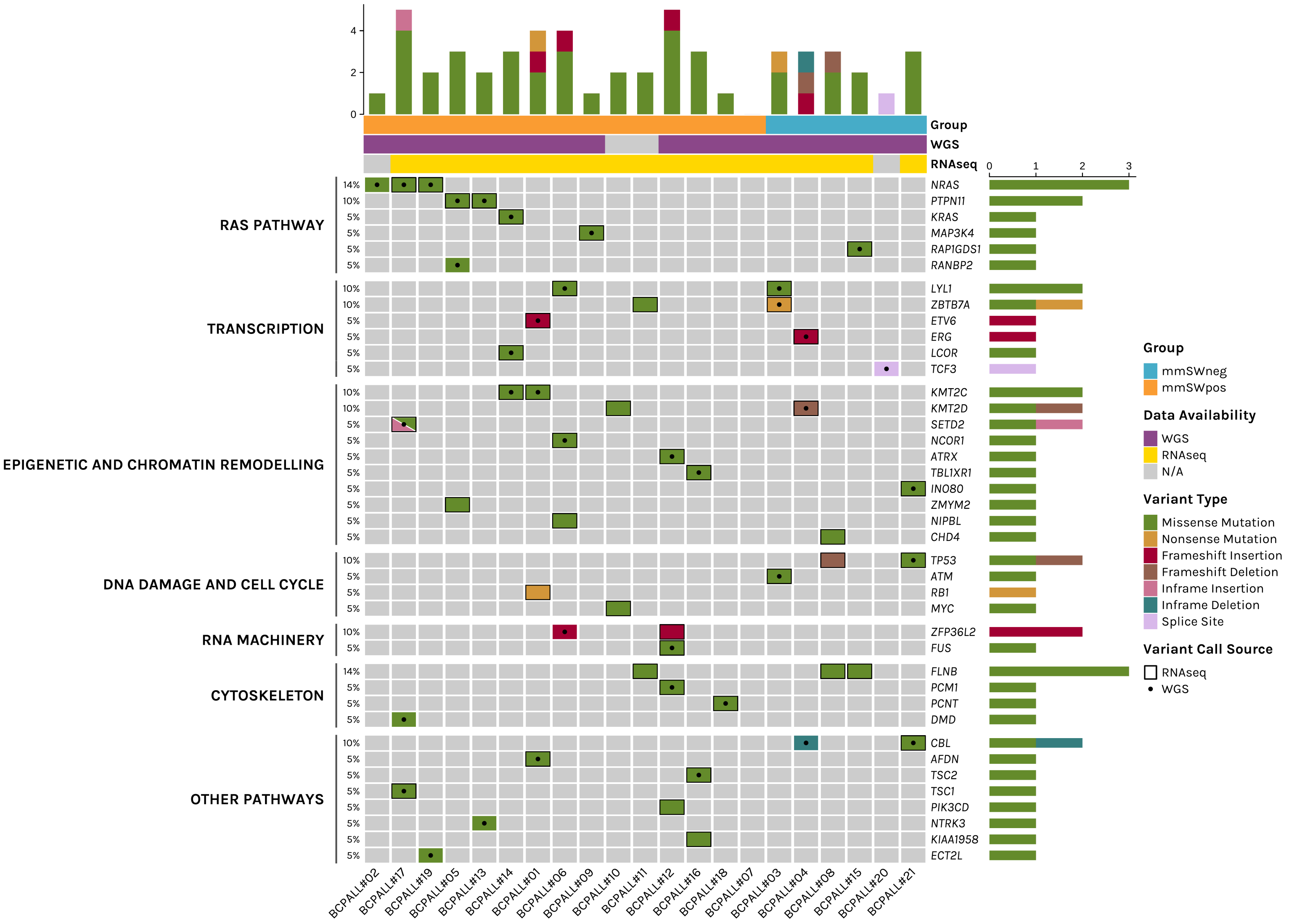


Oncoplot showing variants identified in BCP-ALL patients at Dx by WGS and RNA-seq (n = 6 mmSWneg and n = 15 mmSWpos). MmSWneg are reported in blue, while mmSWpos in orange. Purple and yellow indicate the source of data, being WGS and RNA-seq, respectively. Variant types are reported in different colors, based on their classification. In each cell, a bold black border and round circle indicates that the variant was identified on RNA-seq and WGS data, respectively.
